# Reliability, sensitivity and predictive value of fMRI during multiple object tracking as a marker of cognitive training gain in combination with tDCS in stroke survivors

**DOI:** 10.1101/603985

**Authors:** Knut K. Kolskår, Geneviève Richard, Dag Alnæs, Erlend S. Dørum, Anne-Marthe Sanders, Kristine M. Ulrichsen, Jennifer Monereo Sánchez, Hege Ihle-Hansen, Jan E. Nordvik, Lars T. Westlye

## Abstract

Computerized cognitive training (CCT) combined with transcranial direct current stimulation (tDCS) has showed some promise in alleviating cognitive impairments in patients with brain disorders, but the robustness and possible mechanisms are unclear. In this prospective double-blind randomized clinical trial, we investigated the feasibility and effectiveness of combining CCT and tDCS, and tested the predictive value of and training-related changes in fMRI-based brain activation during attentive performance (multiple object tracking) obtained at inclusion, before initiating training, and after the three-weeks intervention in chronic stroke patients (> 6 months since hospital admission). Patients were randomized to one of two groups, receiving CCT and either (1) tDCS targeting left dorsolateral prefrontal cortex (1 mA), or (2) sham tDCS, with 40s active stimulation (1 mA) before fade out of the current. 77 patients were enrolled in the study, 54 completed the cognitive training, and 48 completed all training and MRI sessions. We found significant improvement in performance across all trained tasks, but no additional gain of tDCS. fMRI-based brain activation showed high reliability, and higher cognitive performance was associated with increased tracking-related activation in the dorsal attention network (DAN) and default mode network (DMN) as well as anterior cingulate after compared to before the intervention. We found no significant associations between cognitive gain and brain activation measured before training or in the difference in activation after intervention. Combined, these results show significant training effects on trained cognitive tasks in stroke survivors, with no clear evidence of additional gain of concurrent tDCS.

## Introduction

While stroke mortality has decreased during the last decades in western countries, it remains a leading cause of disabilities (Feigin et al., 2017), and many stroke survivors experience persistent cognitive sequelae. Indeed, prevalence of neurocognitive disorders in post stroke survivors have been reported to as high as 54% (Barbay et al., 2018), and may increase risk of dementia (Pendlebury, 2012). Reduced working memory capacity is among the most common impairments (Teasell, 1992; Zinn et al., 2007) with high prevalence at the chronic stage (Mahon et al., 2017; Schaapsmeerders et al., 2013), and potentially large impact on daily functioning (Synhaeve et al., 2014). Stroke lesions influence brain activation measured by brain imaging (Altamura et al., 2009), and working memory deficits have been linked to aberrant brain activation during task engagement (Ziemus et al., 2007).

Computerized cognitive training (CCT) has shown some promise in augmenting cognitive rehabilitation. A recent meta-analysis revealed small but significant improvements on cognitive functioning in response to CCT in healthy participants (Au et al., 2015). Similar findings have been reported in patients with Parkinson’s disease (Leung et al., 2015), mild cognitive impairment (Maffei et al., 2017), brain injury (Weicker et al., 2016), and stroke (De Luca et al., 2018b; Peers et al., 2018), as well as schizophrenia (Prikken et al., 2019) and major depression (Motter et al., 2016). However, the underlying mechanisms remain elusive.

Furthermore, non-invasive electrical stimulation of the brain has been suggested to augment training benefits (Kang et al., 2016; Marquez et al., 2015). Indeed, evidence suggests amplified motor recovery post-stroke, when physical therapy is paired with transcranial direct current stimulation (tDCS) (Kang et al., 2016), and the beneficial effects have been suggested to generalize to a cognitive rehabilitation context (Andrews et al., 2011; Fregni et al., 2005; Lawrence et al., 2018). Despite some positive findings, the overall effectiveness of tDCS has been questioned (Medina and Cason, 2017). Also, the potential mechanisms, if any, are unclear, yet enhancing neuronal excitability by lowering the membrane potential is one of the suggested mechanisms (Giordano et al., 2017).

Functional MRI has become an increasingly used non-invasive brain imaging method linking behavior to brain activation patterns. Studies investigating alterations in brain activation in response to CCT have reported mixed results. In healthy controls, training gain has been associated with decreased activity in medial and dorsal frontal areas, including anterior cingulate (Heinzel et al., 2016; Miró-Padilla et al., 2019; Motes et al., 2018), with increased efficiency reflected in lowered activity as a suggested cognitive mechanism (Constantinidis and Klingberg, 2016). For patient cohorts the current evidence is sparser, but increased activity in frontal and parietal areas in response to cognitive training has been reported in patients with schizophrenia (Haut et al., 2010; Ramsay and MacDonald, 2015; Subramaniam et al., 2014). In stroke patients, CCT has been associated with increased functional connectivity (FC) between hippocampus, frontal and parietal areas, which was associated with improved cognitive performance (Lin et al., 2014; Yang et al., 2014).

Despite some promising findings, the effects of CCT (Buitenweg et al., 2019; De Luca et al., 2018a; Melby-Lervåg and Hulme, 2013; Melby-Lervåg et al., 2016) and tDCS (Dedoncker et al., 2016; Hill et al., 2016) are still unclear. More knowledge about the origins of individual differences in training gains is relevant to evaluate and improve selection to fully realize the clinical potential of these interventions. Investigations of the neural underpinnings and mechanisms of cognitive improvement in response to CCT and tDCS may help clarify the benefits and limitations for future clinical application.

To this end, we investigated differences in training-related gains of a commonly used working memory computerized training program in combination with tDCS, and tested the predictive value and sensitivity of fMRI-based brain activation as a marker for training-related gain in task performance. Specifically, we estimated activation patterns during multiple object tracking (MOT) at baseline, before initiating training (on average 4 weeks after baseline measure), and after a three-week intervention. The multiple object tracking (MOT) task offers a promising tool for assessing and manipulate converging activation patterns across attention and working memory tasks in both healthy controls and stroke patients (Dørum et al., 2016; Pylyshyn and Strom, 1988; Walle et al., 2019). MOT robustly recruit brain areas (Alnæs et al., 2014) comparable with activations associated with Cogmed training (Olesen et al., 2004), including canonical task-positive and task-negative brain networks. To assess specific associations with tDCS and following a double-blind design, in addition to CCT each participant was pseudo-randomized into two groups: 50% (n=27) receiving active tDCS directed towards the left dorsolateral prefrontal cortex and 50% (n=27) receiving sham stimulation.

We tested the following hypotheses: (1) Variability in cognitive training gain is reflected in differential slopes of increased performance across participants, (2) level of improvement is affected by tDCS stimulation protocol, (3) fMRI-based brain activation patterns prior to training are predictive of training gain, and (4) individual differences in task improvement are reflected in differential patterns of brain activation after the three-weeks intervention, in particular involving frontal and parietal brain regions.

## Methods

### Sample and exclusion criteria

Figure 1 depicts the study recruitment pipeline. Participants were recruited from two major hospitals located in Oslo, Norway (Stroke unit at Oslo University hospital, and Geriatric department at Diakonhjemmet Hospital, Oslo, Norway). Hospital staff identified suitable participants. Inclusion criterion was identifiable stroke with ischemic or hemorrhagic etiology. Patients with transient ischemic attacks (TIA) were not included. Exclusion criteria included MRI contraindications, other neurological diseases diagnosed prior to the stroke, severe psychiatric diagnosis including bipolar disorder, schizophrenia or drug abuse. Approximately 900 invitations were sent out, and 250 responded; among those, 77 patients were eligible for participation and were initially included in the study (Ulrichsen et al., 2020). 19 participants withdrew from the study prior to (n=14) or during (n=5) the cognitive training due to the labor-intensity of the intervention. Four participants were excluded during the course of the intervention due to newly occurring medical issues violating the inclusion criteria. Two participants were included without radiological findings when admitted to the stroke unit, but displayed neurological symptoms. Initial MRI diffusion weighted imaging revealed possible acute infarctions and the patients were treated with thrombolysis. Subsequent T2-weighted imaging revealed no certain radiological findings, but symptoms persisted upon discharge and they were diagnosed by a neurologist with ICD-10 code I63.9 Cerebral infarction, and therefore included. In total, 54 participants completed the cognitive training program. For timepoint 1, one participant had corrupted behavioral data and two participants were not able to complete the task during the fMRI assessment, and were excluded from all MRI analysis. For MRI timepoint 2 and 3, one participant had poor alignment to the MNI-template, and two participants were not able to complete the task, yielding a total of 51 participants at baseline and 48 with full MRI at all timepoints. In this final sample, 26 participants received active stimulation, and 22 received sham stimulation.

**Figure 1.**
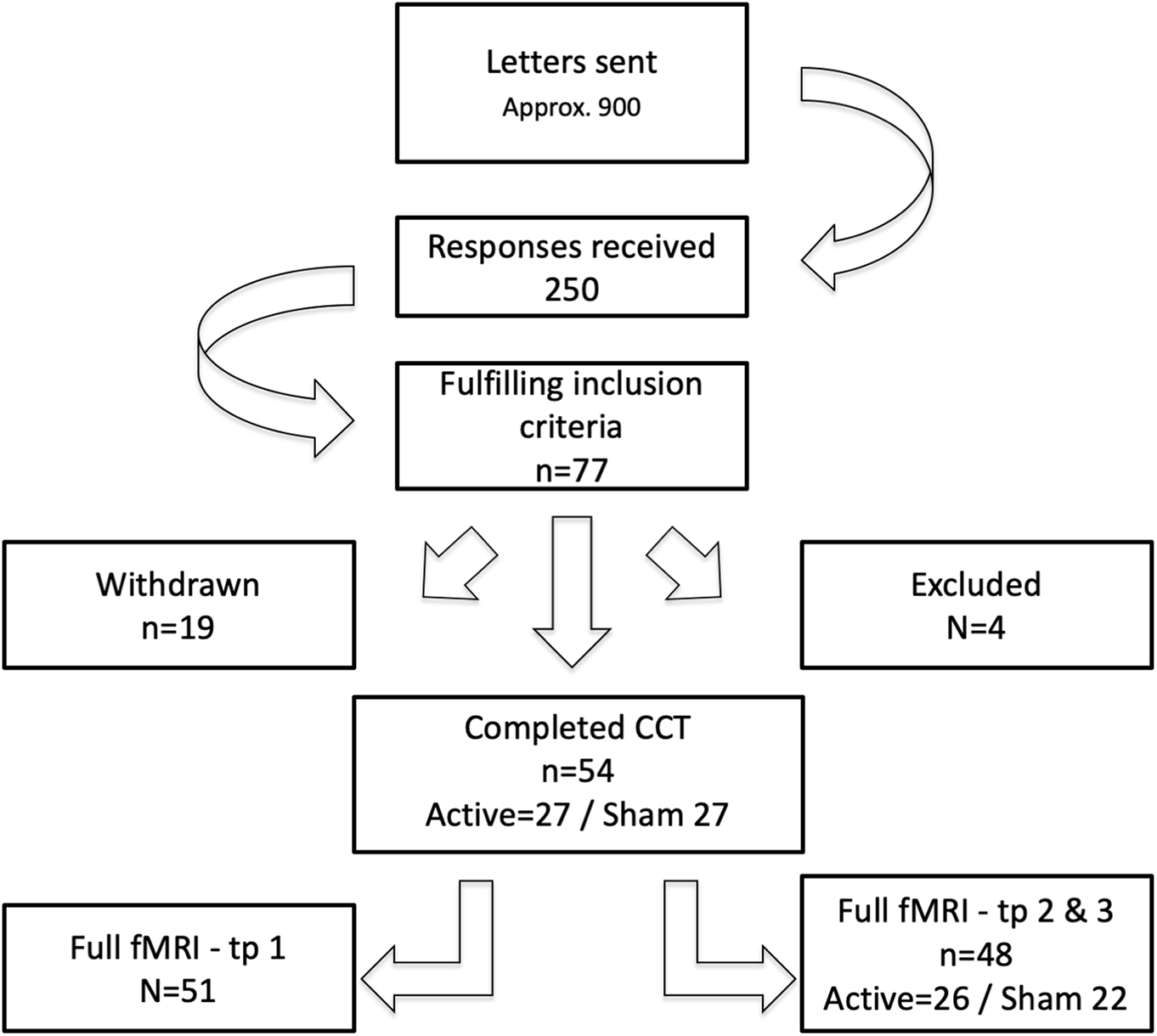
Flowchart displaying inclusion pipeline

The study was approved by the Regional Committee for Medical and Health Research Ethics South-East Norway (2014/694; 2015/1282), and all participants provided written informed consent.

### Patient characteristics and lesion demarcation

Sample characteristics for the participants completing the intervention (n=54) are described in Table 1, including National Institutes of Health Stroke Scale (NIHSS) (Goldstein et al., 1989), Trial of Org 10172 in Acute Stroke Treatment (TOAST) (Adams et al., 1993), Mini-Mental State Exam (MMSE) (Strobel and Engedal, 2008) and IQ derived from Wechsler Abbreviated Scale of Intelligence (WASI) (Ørbeck and Sundet, 2007). For an extended overview of the separate intervention groups, see supplementary table 1. Briefly, the sample mean age was 69 years (sd=7.3), comprised 74% males, and average IQ of 110 (sd=16). Two sample t-tests revealed no significant differences between the active and sham tDCS-groups.

**Table 1.** Sample characteristics. Two-sample t-tests did not reveal any significant differences between the active and sham group regarding participant characteristics

|  | Mean | sd | Min | Max | tDCS<br>grouping<br>t/p |
| --- | --- | --- | --- | --- | --- |
| **Participant descriptives (current)** |  |  |  |  |  |
| Age at inclusion | 69.13 | 7.374 | 47 | 81 | 0.11 / 0.907 |
| Males (%) | 74.07 | - |  |  | 23 / 17 males |
| Time between stroke and inclusion (months) | 25.74 | 9.17 | 6 | 45 | 0.36 / 0.720 |
| Self-reported education in years | 14.39 | 3.71 | 9 | 30 | -0.59 / 0.559 |
| MMSE at inclusion | 27.98 | 1.855 | 22 | 30 | 0.81 / 0.419 |
| IQ | 110.45 | 16.906 | 66 | 136 | -0.41 / 0.687 |
| Days between inclusion and training | 33.78 | 12.986 | 19 | 74 | 0.82 / 0.419 |
| Days between MRI assessment pre/post training | 31.72 | 6.257 | 19 | 49 | 1.99 / 0.052 |
| **Clinical variables during hospitalization** |  |  |  |  |  |
| NIHSS at discharge | 1.33 | 1.53 | 0 | 7 |  |
| TOAST classification | Large vessel disease n=20<br>Small vessel disease n=18<br>Cardioembolic disease n=6<br>Other n=10 |  |  |  |  |

For each participant, lesion demarcation was performed by a trained assistant, utilizing the semi-automated toolbox clusterize, implemented for SPM8 (Clas et al., 2012; de Haan et al., 2015), and guided by radiological descriptions. Figure 2 displays a probabilistic lesion map across participants. Supplementary Figure 1 provides an overview of individual lesions.

**Figure 2.**
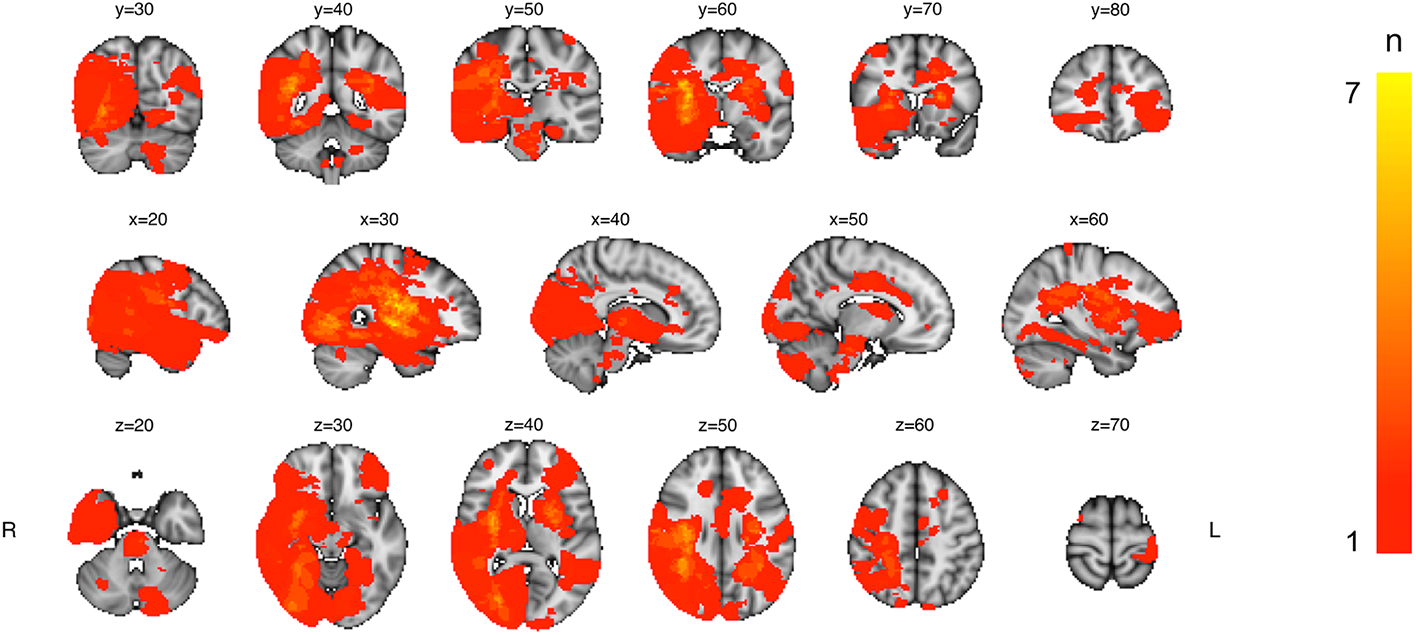
Heatmap displaying lesion overlap across included participants. The color scale reflects the number of overlapping lesions in each voxel (ranging from 0 to 7).

### Study protocol

The protocol has previously been described in detail (Richard et al., 2020). Briefly, after inclusion the participants completed MRI assessment at three timepoints: timepoint 1, at timepoint 2 on average 3-4 weeks after timepoint 1 without additional contact to enable a double baseline for comparison, and at timepoint 3, after the participant completed the training protocol.

### Cognitive training

The patients performed a computerized working memory training program (Cogmed QM, Cogmed Systems AB, Stockholm, Sweden), comprising 12 auditory-verbal and visual-spatial working memory tasks. The implementation of the protocol has previously been reported in detail (Richard et al., 2020). Briefly, the training comprised five weekly training sessions over 3-4 weeks, with 17 training sessions for each participant in total. Each participant completed two weekly tDCS stimulation sessions at the hospital, with a total of 6 tDCS sessions with a minimum of 48 hours between each stimulation. The remaining Cogmed sessions was completed at home. Level of Cogmed task difficulty was automatically adjusted according to each participant’s performance level. The initial two first sessions for each task was discarded from statistical analysis to ensure each participant was at a desired level of difficulty. The same task setup was used for all participants, and tasks completed three times or less over the course of training were excluded. Eight Cogmed subtests were included in further analysis, including Grid, Cube, Digits, Hidden objects, Sort, Rotation, Twist, and 3D-cube.

### tDCS

Participants were randomly assigned to either active or sham condition, using codes provided by the manufacturer and implemented by an in-house Matlab script, pseudo-randomizing participants while maintaining balance in groups of 20. tDCS stimulation was applied at six occasions for each participant, with a minimum of 48 hours between stimulations, aiming at an average of two stimulations per week. Stimulation current was 1000 μA, stimulation time was 20 minutes for the active group with a ramp-up time of 120 seconds and fade-out time of 30s. The current intensity was limited to 1mA to limit the risk of adverse events due to the electrical stimulation. The sham stimulation consisted of ramp-up followed by 40 seconds of active stimulation and then fade-out, following factory settings. Stimulation was delivered via a battery-driven direct current stimulator (neuroConn DC stimulator plus, Germany), through 5 × 7 cm rubber pads, yielding a 28.57 μA/cm2 current density at skin surface. Impedance threshold at start-up of stimulation was < 20 kΩ, and was regulated by the device with an absolute threshold at 40 kΩ for automatically terminating stimulation. The pads were covered with high-conductive gel (Abralyt HiCl, Falk Minow Services Herrsching, Germany) to enhance conductance.

Following previous findings suggesting stimulation of the left dlPFC is associated with working memory improvement when paired with relevant tasks (Andrews et al., 2011; Mancuso et al., 2016), the anodal electrode was placed over F3 corresponding to left DLPFC and cathodal at O2 corresponding to right occipital/cerebellum following the 10-20 system, and fixated with rubber bands.

### fMRI task

Participants performed a blocked version of the MOT (Pylyshyn and Strom, 1988), as previously described in detail (Alnæs et al., 2014). Briefly, the participants were presented with ten identical circles on a grey background. All objects were initially blue, before zero (passive viewing), one (load 1) or two (load 2) of the objects turned red for 2.5 seconds, designating them as targets. After targets had returned to blue, the objects started moving randomly around the screen for 12 seconds. After the 12 second tracking period, the objects became stationary, one of the objects turned green, and the participants were instructed to respond “yes” or “no” to whether or not the green probe had been designated as a target in that trial. Participants were instructed to keep fixation on a fixation point at the center of the screen during both attentive tracking and passive viewing. The task consisted of 24 semi-randomized trials in total, ensuring the same condition was not repeated consecutively.

### MRI acquisition

MRI data was acquired with a 3T GE 750 Discovery MRI scanner (32-channel head coil) located at Oslo University Hospital. T1-weighted images was acquired using a 3D IR-prepared FSPGR (BRAVO) with the following parameters: scan time: 4:43 minutes and included 188 sagittal slices, repetition time (TR): 8.16 ms, echo time: 3.18 ms, flip angle: 12°, voxel size: 1 × 1 × 1 mm, field of view: 256 × 256 mm,

*Functional MRI sequences* was acquired with a BOLD-sensitive gradient echo planar sequence, with TR of 2.25 s, TE 30 ms, field of view 256 × 256 mm, slice thickness 3 mm with 0.5 mm gap acquired ascending interleaved, 90° flip angle. The initial first five volumes were removed before analysis.

### fMRI data processing

fMRI data was processed using FMRI Expert Analysis Tool (FEAT) Version 6.00, from FMRIB’s Software Library (FSL)(Jenkinson et al., 2012; Smith et al., 2004), and included the following steps: correction for motion using MCFLIRT (Jenkinson et al., 2002), linear trend removal and high-pass filtering (0.01 Hz), removal of non-brain tissue using BET (Smith, 2002), spatial smoothing with a Gaussian kernel of full width at half maximum (FWHM) of 6 mm (Smith and Brady, 1997), and linear registration using FLIRT (Jenkinson and Smith, 2001) to Montreal Neurological Institute (MNI) 152 standard space using the T1-weighted scan as an intermediate. In the same manner, normalization parameters were estimated for each participants T2-weighted image, and applied to the demarked lesion.

### Statistical analysis of behavioral data

#### Cogmed main effects of time

To investigate the trajectory of Cogmed performance across participants and time points, linear mixed effects (lme) models were estimated for each test, using the lme function from the nlme-package (Pinheiro et al., 2017) in R (Team, 2016). Task performance was entered as dependent variable, with training session as independent variable, and age, sex, tDCS-group, educational level in years (self-reported), and interaction between training session and tDCS, as fixed factors, and participant as random factor. We used Cogmed’s estimate of current working memory capacity as a measure of task performance. These individual level estimates are derived using an adaptive algorithm, in which task difficulty is increased or decreased in response to correct and incorrect responses. To assess normality according to criteria for lme, we plotted distribution of random effects intercepts, see supplementary Figure 7. Visual inspection did not indicate severe violations of the criteria.

#### Cogmed individual trajectories

To quantify individual trajectories in Cogmed performance across the training-period, we estimated for each test and each participant a linear model with performance as dependent variable and session number as independent variable, yielding a beta-estimate (slope) for each test for each participant reflecting changes in performance over time. As a measure of overall performance for each participant, we used the average task performance across sessions for each test for each participant. To investigate task homogeneity in beta-estimates and average performance we performed multivariate outlier detection with the aq.plot function in the mvoutliers package in R (Filzmoser and Gschwandtner, 2018), with an alpha of 0.001 and 90% of the sample to estimate the minimum covariance determinant. Two subtests (“hidden objects” and “Digits”) were found to have a high number of outliers compared to the other subtests, and were excluded from further analysis (See Supplementary Figure 2 for details).

To derive a common score for average Cogmed performance and gain, we performed a principal component analysis (PCA) on beta-estimates and average performance, respectively. All scores were zero-centered and standardized prior to running the PCA. We used the first factor from the beta PCA as a *Cogmed change score*, and the first factor from the average performance PCA as a *Cogmed average performance score (*see Supplementary Figure 3 for scree-plot derived from the PCA performed on both with and without the excluded tasks).

To explore potential association between initial cognitive capacity and Cogmed change score, we correlated performance on the cognitive performance at baseline assessed with the Cabpad battery (https://www.cognisoft.info) at TP1, described in detail previously (Richard et al., 2020) with the Cogmed change score. To test for associations between initial cognitive performance and response to tDCS we estimated linear models using the same baseline cognitive performance from Cabpad as the dependent variable, and Cogmed change score and tDCS group as independent variables. Models were estimated using the lm function in the stats package in R (Team, 2016).

To assess the association between Cogmed scores and lesion severity, we estimated linear models for both Cogmed change and average performance scores, using lesion volume, number of lesions and their interaction as explanatory variables. To assess impact of duration from the stroke to inclusion on gains from the training, we correlated time since injury measured in months with the Cogmed change score as well as the Cogmed average performance score.

#### fMRI behavioral analysis

To investigate performance during the MOT task across timepoints, linear mixed effects (lme) models were estimated for both load 1 and load 2, using the lme function from the nlme-package (Pinheiro et al., 2017) in R. Task performance was entered as dependent variable, with training session as independent variable, and participant as random factor. To assess for pairwise differences across the timepoints, post hoc analysis was performed using the ghlt function from the multcomp package (Hothorn et al., 2020) in R, and corrected for multiple comparisons using Bonferroni correction.

### fMRI data analysis

First-level GLMs for each participant and scanning session were estimated using FEAT (Woolrich et al., 2001), including regressors for passive viewing, load 1, load 2, responses, as well as the standard and extended motion parameters estimated during preprocessing. The following contrasts were estimated: passive viewing, load 1, load 2, tracking ([load 1 + load 2] – passive viewing) and load (load 2 – load 1).

To estimate main effects of experimental conditions across individuals, contrasts of parameter estimates (COPEs) from each participant’s first scan session (baseline) were submitted to randomise (Winkler et al., 2014) performing a one sample t-test for each condition separately. Multiple comparisons were corrected for using permutation testing and threshold free cluster enhancement (TFCE (Smith and Nichols, 2009)).

To estimate associations with task performance with brain activation during task engagement, individual COPE maps for load 1 and load 2 were stacked separately, and submitted to randomise, including sex, age and task performance as covariates.

To test for differences in brain activation between scan sessions, we first performed fixed effects analysis for each participant separately, including all COPEs from first level GLM, specifying the contrasts session 3 minus session 2, session 3 minus session 1, and session 2 minus session 1. Next, all individual COPE maps were submitted to randomise to test for group-level main effects while correcting for multiple comparisons using permutation testing and threshold free cluster enhancement (TFCE).

To quantify within-subject reliability in brain activation across time we estimated intraclass correlation coefficient (ICC) for each voxel between assessment two and three utilizing the variance from one-way random ANOVA, for the contrasts passive viewing, load 1 and load 2, as implemented in the ICCest function from the ICC R package (Wolak et al., 2012).

To test for associations between Cogmed change and average performance scores and brain activation, individual level COPE maps for timepoint 1 and the difference between timepoint 3 and 2 including the conditions passive viewing, load1, load 2, tracking as well as load were stacked separately and submitted to randomise. The model included sex, age, and either Cogmed change score or Cogmed average performance score (obtained from the PCA). In the same manner to explore differences associated with tDCS stimulation protocol, the same COPE’s from the difference between timepoint 3 and 2 including all conditions were submitted to randomise, with the model including sex, age and tDCS-condition. All models were corrected for multiple comparisons across space using 5000 permutations and thresholded using TFCE.

## Results

### Cogmed performance

Figure 3 shows Cogmed performance over time for each group and Table 2 displays corresponding summary statistics. Lme revealed a significant group-level improvement on all Cogmed tests included in the analysis. In addition, we found a significant association with age on mean performance across all tests, indicating lower average performance with higher age. We found no significant associations between Cogmed performance and sex, tDCS group (sham/experimental), or years of education. We found a significant interaction between time and tDCS on the twist task, suggesting augmented training gain in the participants in the active tDCS stimulation group. However, after removal of one outlier defined by visual inspection, the result did not reach significance (see Supplementary Table 2 and Supplementary Figure 4 for corresponding statistics and visualization).

**Figure 3.**
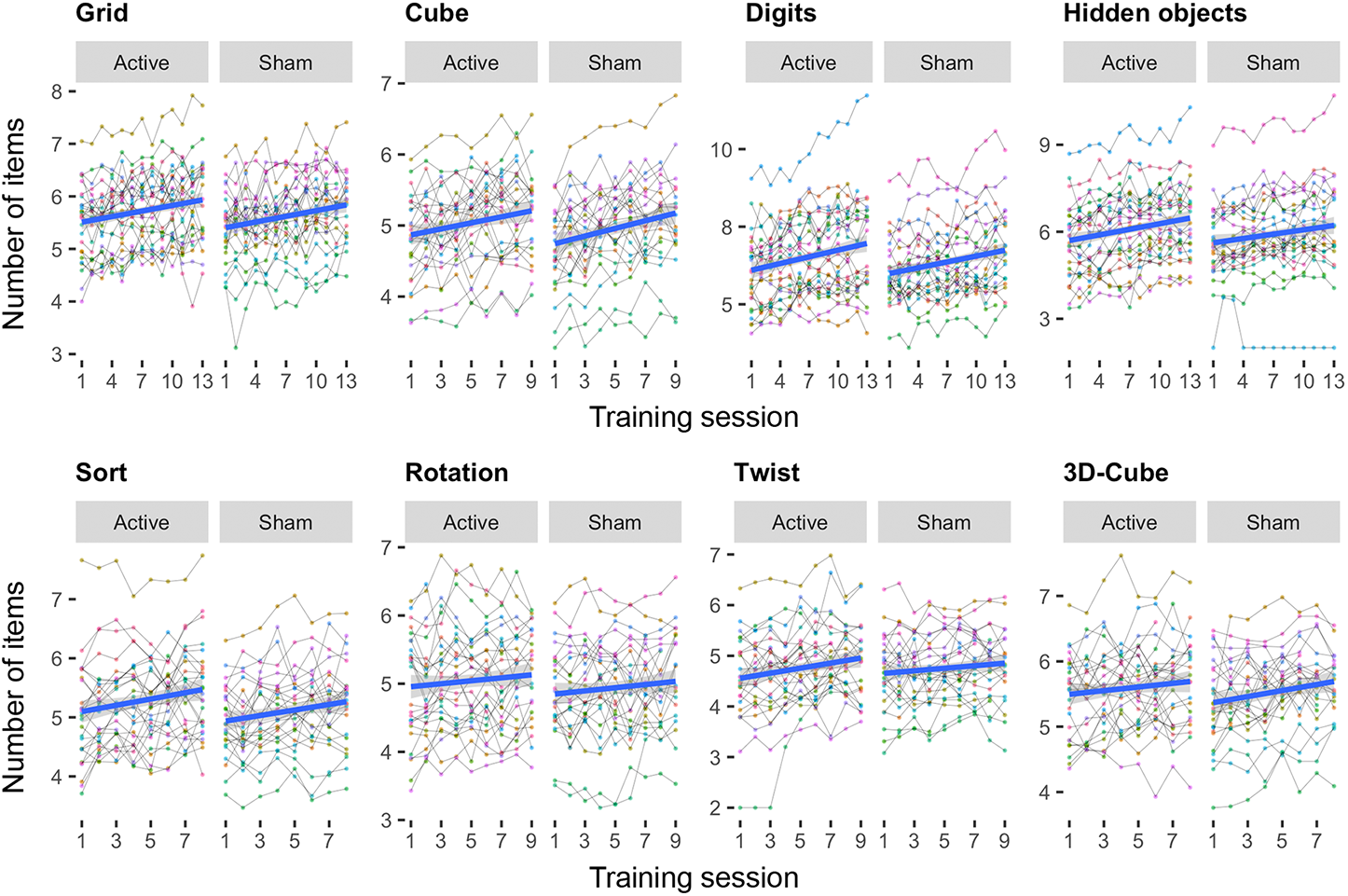
Individual level task performance during the course of the intervention period for each group (tDCS sham/active). The fit lines are based on a linear fit within each group.

**Table 2.** Summary statistics (t/(p))from linear mixed effects models testing for associations between task performance and session, age, sex, tDCS, education and the interaction between session and tDCS. t-vaules along with raw p-vaules are displayed, significant findings are highlighted (FDR-corrected, alpha = 0.05).

|  | Grid | Sort | Digits | Cube | Hidden objects | Twist | 3D-Cube | Rotation |
| --- | --- | --- | --- | --- | --- | --- | --- | --- |
| Session | **8.66**<br>(*<0.001*) | **6.40**<br>(*<0.001*) | **12.79**<br>(*<0.001*) | **7.44**<br>(*<0.001*) | **11.03**<br>(*<0.001*) | **6.45**<br>(*<0.001*) | **3.16**<br>(*0.002*) | **2.21**<br>(*0.028*) |
| Age | **-3.05**<br>(*0.004*) | **-2.11**<br>(*0.041*) | **-6.07**<br>(*<0.001*) | **-2.54**<br>(*0.015*) | **-4.86**<br>(*<0.001*) | **-3.62**<br>(*0.001*) | **-3.90**<br>(*<0.001*) | **-3.41**<br>(*0.001*) |
| Sex | -0.89<br>(0.377) | -0.33<br>(0.741) | -2.29<br>(0.027) | -1.22<br>(0.23) | -2.25<br>(0.029) | -1.46<br>(0.15) | -1.42<br>(0.162) | -1.34<br>(0.187) |
| tDCS | -0.65<br>(0.521) | -1.03<br>(0.309) | -0.06<br>(0.951) | -0.53<br>(0.599) | 0.12<br>(0.908) | 1.00<br>(0.324) | -0.82<br>(0.414) | -0.81<br>(0.422) |
| Education | 0.06<br>(0.951) | 0.96<br>(0.34) | 0.90<br>(0.374) | 0.14<br>(0.886) | 1.57<br>(0.124) | 0.01<br>(0.991) | -0.15<br>(0.884) | 0.33<br>(0.744) |
| Session x tDCS | 0.2<br>(0.842) | -0.39<br>(0.7) | -1.53<br>(0.128) | 0.46<br>(0.644) | -1.79<br>(0.074) | **-2.83**<br>(*0.005*) | 1.72<br>(0.087) | 1.23<br>(0.22) |

We found no significant correlation between Cogmed average performance and Cogmed change score (r=.17, t=1.29, p=.2). We found a significant association between lesion volume and Cogmed change score (t −2.5, p=.017), indicating a negative correlation between lesion volume and training response, which did not retain significance after removal of one outlier (t=−.58, p=.563). See Supplementary Table 3 for corresponding statistics. Furthermore, results revealed no significant correlations between time since injury and Cogmed average performance (r= −0.19, t = −1.38, p = 0.173) nor Cogmed change score (−0.19, t = −1.35, p = 0.181).

Linear models revealed no significant associations between cognitive capacity as measured using Cabpad at baseline and gain in Cogmed performance. Furthermore, there were no significant interactions between Cabpad score at baseline and tDCS on Cogmed change score. See supplementary Figure 6 and supplementary table 5 for corresponding statistics.

### MOT task performance

LME analysis revealed main effect of time on task performance during both load 1 and load 2. Post hoc analysis revealed significant differences between timepoint 3 and 1, as well as between timepoint 2 and 1, but not between timepoint 3 and 2. See Table 3 and Figure 4 for details.

**Table 3.**
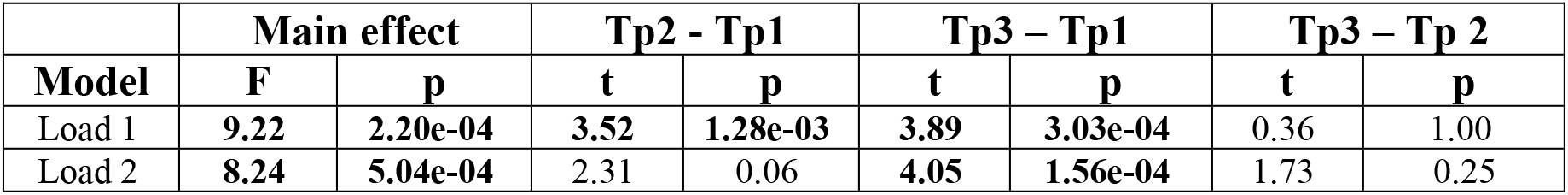
Summary statistics from linear mixed effects models testing for associations between task performance and session for the MOT task. Bonferroni-corrected p-values are displayed, significant results are highlighted.

|  | Main effect |  | Tp2 - Tp1 |  | Tp3 – Tp1 |  | Tp3 – Tp 2 |  |
| --- | --- | --- | --- | --- | --- | --- | --- | --- |
| Model | F | p | t | p | t | p | t | p |
| Load 1 | 9.22 | 2.20e-04 | 3.52 | 1.28e-03 | 3.89 | 3.03e-04 | 0.36 | 1.00 |
| Load 2 | 8.24 | 5.04e-04 | 2.31 | 0.06 | 4.05 | 1.56e-04 | 1.73 | 0.25 |

**Figure 4.**
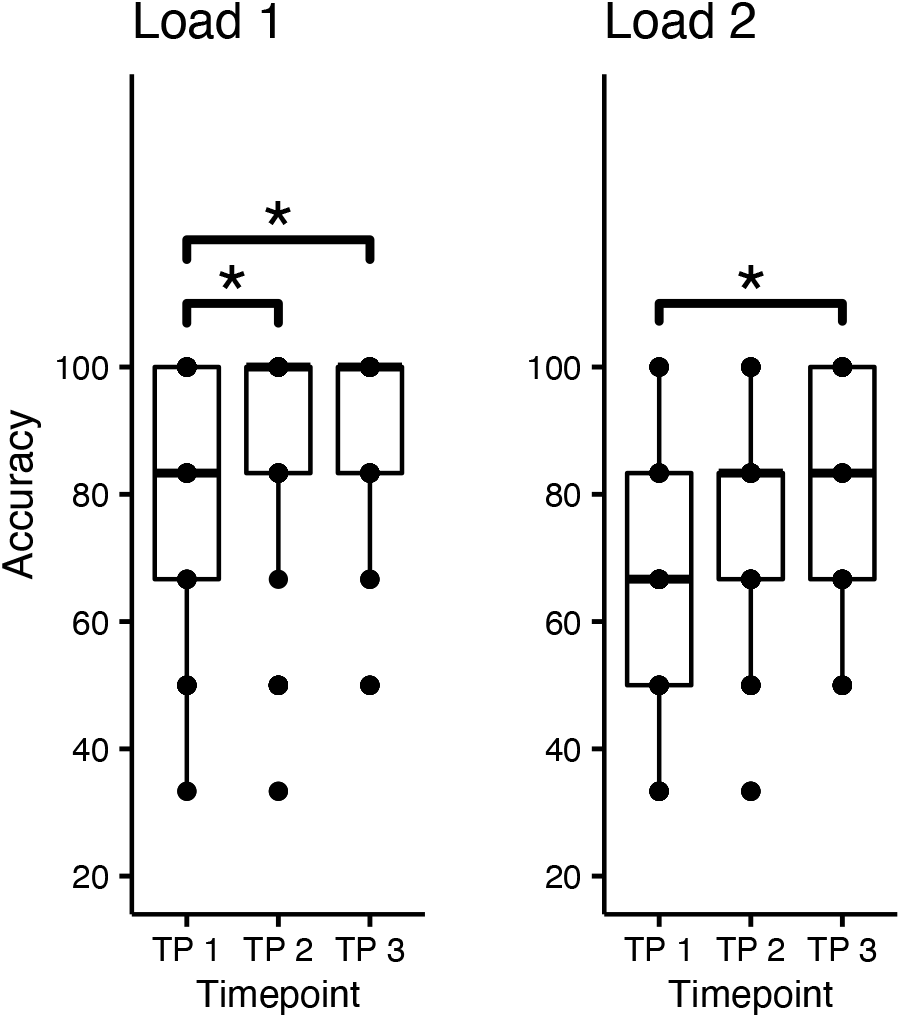
Task performance during the MOT task at the three different timepoints. Significant differences in group means is marked *, Table 3 displays corresponding statistics.

### Association between MOT performance and fMRI activation

Linear models revealed no significant association between in-scanner task performance and fMRI activation during neither load 1 or load 2 during the first assessment (TP 1). In-scanner behavioral performance was therefore not included in further analysis.

### fMRI main effects of task

Figure 5 shows main effects of task condition at baseline, and Table 4 summarizes anatomical locations. Passive viewing was associated with increased activity in the dorsal attention network (DAN) and the inferior and superior occipital cortex, and decreased activation in pre and post-central gyrus, supplementary motor area (SMA), anterior cingulate, as well as default mode network (DMN), including posterior cingulate gyrus (PCC), medial prefrontal gyrus (mPFC), and superior and medial temporal gyrus (S/MTG).

**Figure 5.**
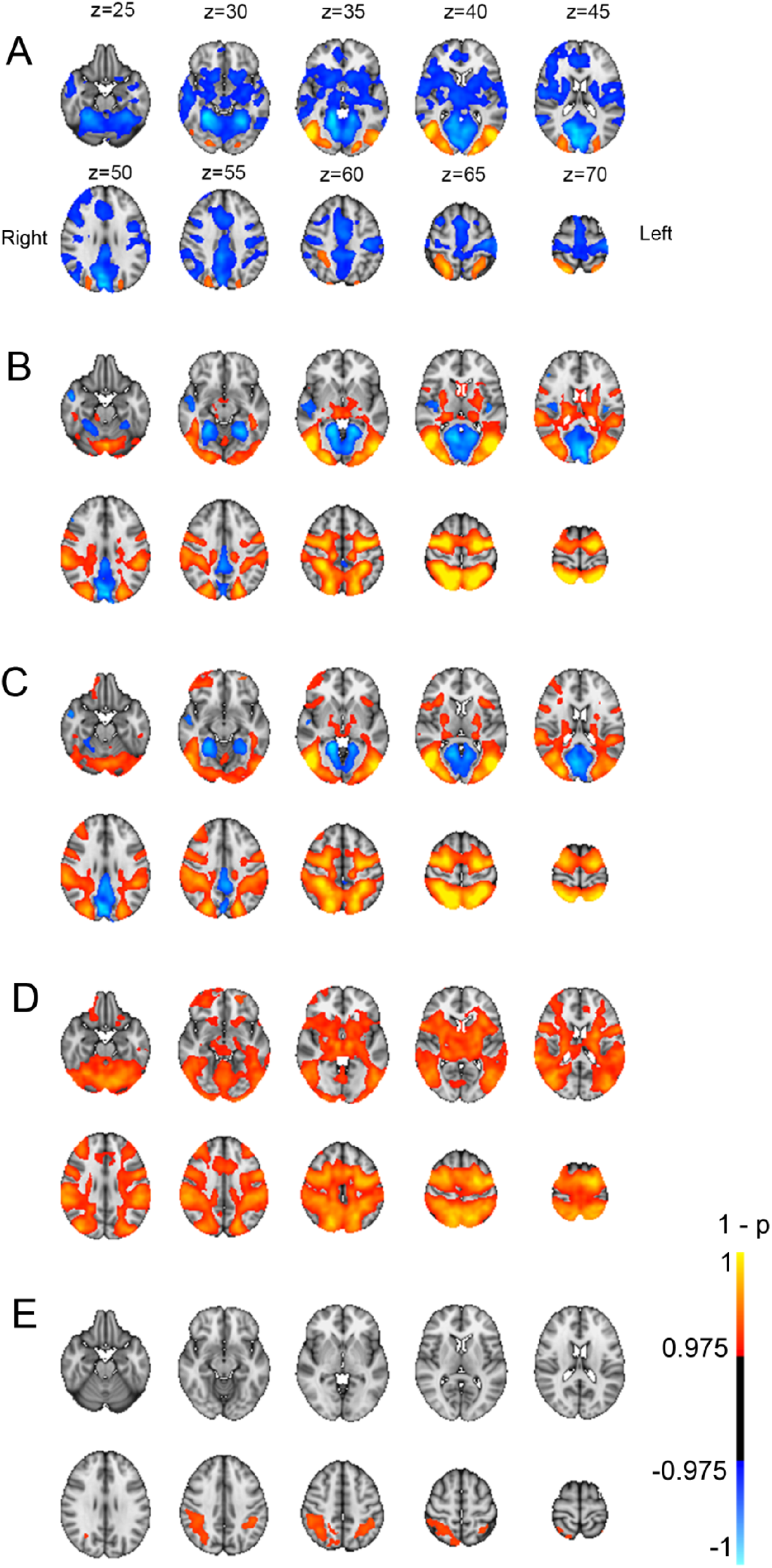
Main effects from task-fMRI for (A) passive viewing > baseline, (B) load 1 > baseline, (C) load 2 > baseline, (D) tracking > passive viewing, and (E) increased cognitive load (load 2 > load 1). Panel A display corresponding coordinates in MNI, and left / right orientation. P-values (1-p) have been multiplied with the sign of the effect in relation to the main contrast.

**Table 4.** Clusters associated with different MOT condition at baseline (all p<.05, corrected)

| Condition | Direction | size | X | Y | Z | Area |
| --- | --- | --- | --- | --- | --- | --- |
| PV | pos | 4029 | 33 | 19 | 30 | Lateral occipital cortex |
|  | pos | 3861 | 55 | 18 | 32 | Lateral occipital cortex |
|  | pos | 1784 | 29 | 41 | 61 | Superior parietal lobe |
|  | neg | 64337 | 33 | 35 | 21 | Cerebellum |
| Load1 | pos | 57023 | 59 | 29 | 8 | Cerebellum |
|  | neg | 12615 | 33 | 39 | 23 | Cerebellum |
|  | neg | 1494 | 19 | 62 | 24 | Middle temporal gyrus |
|  | neg | 256 | 65 | 53 | 42 | Central opercular cortex |
|  | neg | 153 | 20 | 78 | 48 | Middle frontal gyrus |
|  | neg | 19 | 22 | 71 | 50 | Inferior frontal gyrus |
| Load2 | pos | 66242 | 59 | 29 | 8 | Cerebellum |
|  | pos | 411 | 65 | 71 | 37 | Frontal operculum |
|  | pos | 8 | 65 | 80 | 55 | Middle frontal gyrus |
|  | neg | 10269 | 31 | 40 | 26 | Occipital fusiform cortex |
|  | neg | 694 | 18 | 63 | 26 | Superior temporal gyrus |
| Tracking | pos | 101656 | 59 | 29 | 8 | Cerebellum |
| Load | pos | 4590 | 24 | 42 | 55 | Inferior parietal lobe |
|  | pos | 1709 | 67 | 42 | 56 | Inferior parietal lobe |
|  | pos | 245 | 24 | 79 | 54 | Middle frontal gyrus |

Load 1 was associated with increased activity in DAN, extending into the inferior and superior occipital cortex, cerebellum, and precentral gyrus, in the cingulo-opercular (CO) network, including thalamus, frontal operculum and insula, as well as in the ventral attention (VAN) network including temporoparietal junction and supramarginal gyrus, middle frontal gyrus, and frontal operculum. Decreased activity was seen in DMN nodes including PCC, mPFC and S/MTG. Load 2 showed a similar pattern to Load 1, with additional increased activation in right anterior middle frontal gyrus. Tracking over passive viewing revealed same activation pattern as load 1 and load 2, with additional bilateral activation of anterior middle frontal gyrus. Load-related (load 2 –load1) increases in activation were seen in superior and inferior parietal lobe, as well as right anterior middle frontal gyrus, with no significant load-dependent decreases in activation.

### fMRI – reliability

Figure 6 shows voxel-wise ICC for COPE values for passive viewing, load 1 and load 2, between timepoint 2 and 3. The results revealed ICC>0.4 across major portions of the brain, including occipital, temporal, parietal lobe for all conditions, as well as and a large part of the frontal lobes for load 1.

**Figure 6.**
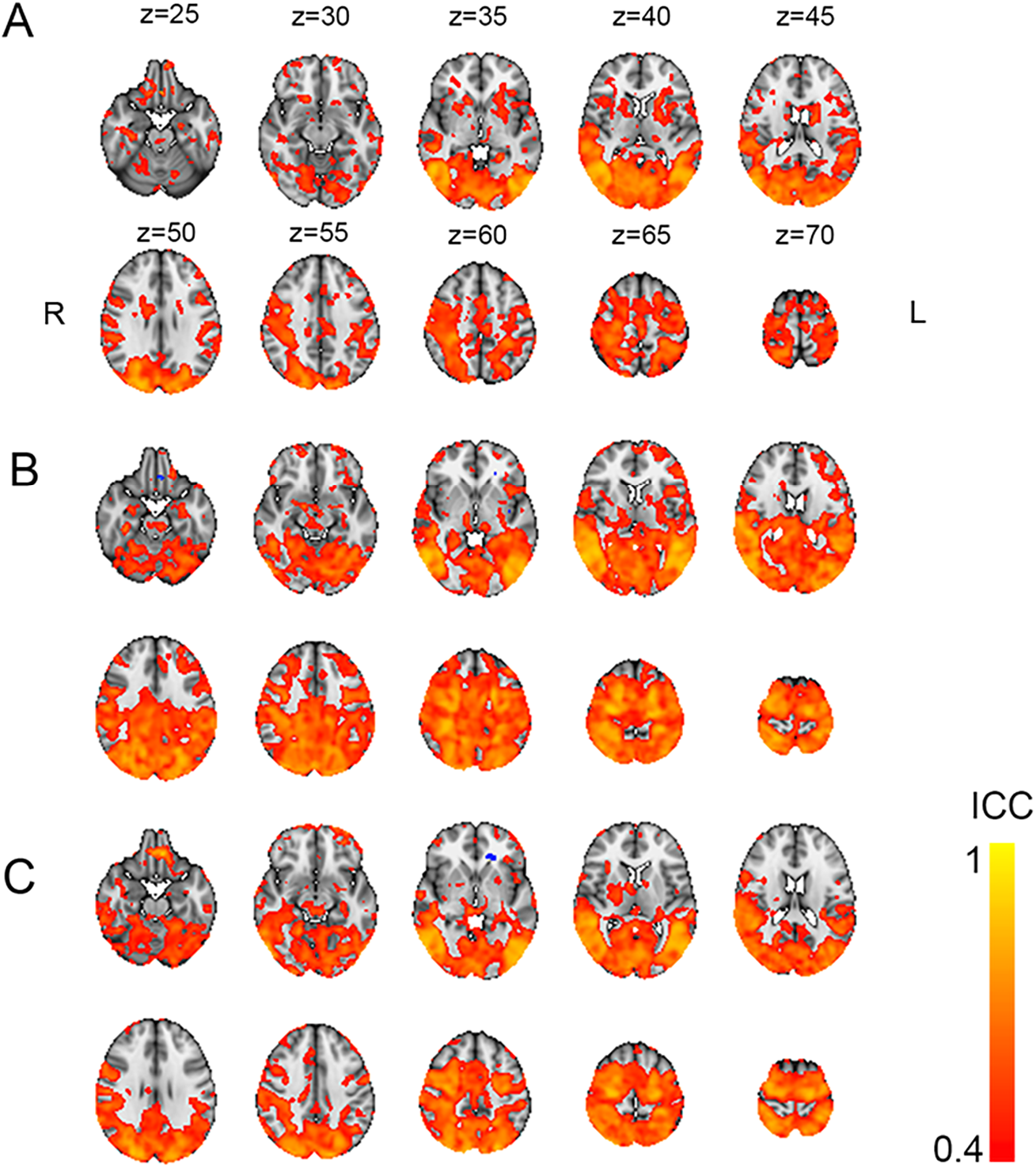
Voxel-wise ICC for condition passive viewing (A), load 1 (B), and load 2 (C). Thresholded at ICC > 0.4.

### Pairwise comparison of fMRI estimates across timepoints

Comparison of fMRI signal between the timepoints revealed no significant differences between timepoint 3 and 1, timepoint 2 and 1 as well timepoint 3 and 2, for neither passive viewing, load 1, load 2, tracking nor load, indicating fluctuating patterns, but not in a time-dependent manner.

### Cogmed –fMRI associations

We found no significant associations between the Cogmed change score and brain activation at timepoint 1, or between Cogmed change score and the difference in brain activation between timepoint 2 and 3.

Furthermore, results revealed no significant associations between the Cogmed average score and brain activation at timepoint 1. However, between timepoint 2 and 3 we found a significant association between Cogmed average performance score and the difference in activation during attentive tracking, indicating larger activation increases in the anterior and posterior parts of the cingulate cortex and left precuneus, superior parietal lobe, lateral occipital cortex, as well as cerebellum with greater average Cogmed performance. See Table 5 and Figure 7 for corresponding statistics. Using a Cogmed average performance score based on all 8 subtests (from the PCA, now including two sub-tests previously excluded), revealed slightly weaker but highly similar effects (See Supplementary Figure 5 and Supplementary Table 4), suggesting that excluding the two subtests did not bias the results.

**Table 5.** Cogmed average score clusters associated with difference in activation between timepoint 2 and 3. (all p<.05, corrected).

|  | Direction | Size | x | y | z | Location |
| --- | --- | --- | --- | --- | --- | --- |
| Frontal | pos | 120 | 30.1 | 64.4 | 61.4 | Middle frontal gyrus |
|  | pos | 46 | 59.5 | 63.9 | 60.5 | MFG L |
| Limbic | pos | 1269 | 41.7 | 74.2 | 53.3 | ACC |
|  | pos | 261 | 43.8 | 52.3 | 53.8 | PCC |
| Parietal | pos | 83 | 23 | 36.8 | 61.9 | SPL |
|  | pos | 207 | 55 | 30.2 | 48.7 | Precuneus cortex |
| Occipital | pos | 117 | 21.5 | 25.9 | 39.3 | Lateral occipital cortex |
|  | pos | 100 | 61.6 | 22.7 | 44.4 | Lateral occipital cortex |
|  | pos | 33 | 35.8 | 15.3 | 36.3 | Occipital pole |
| Cerebellar | pos | 54 | 60 | 26 | 13 | Cerebellum |
|  | pos | 16 | 41.1 | 37.2 | 23.5 | Cerebellum |

**Figure 7.**
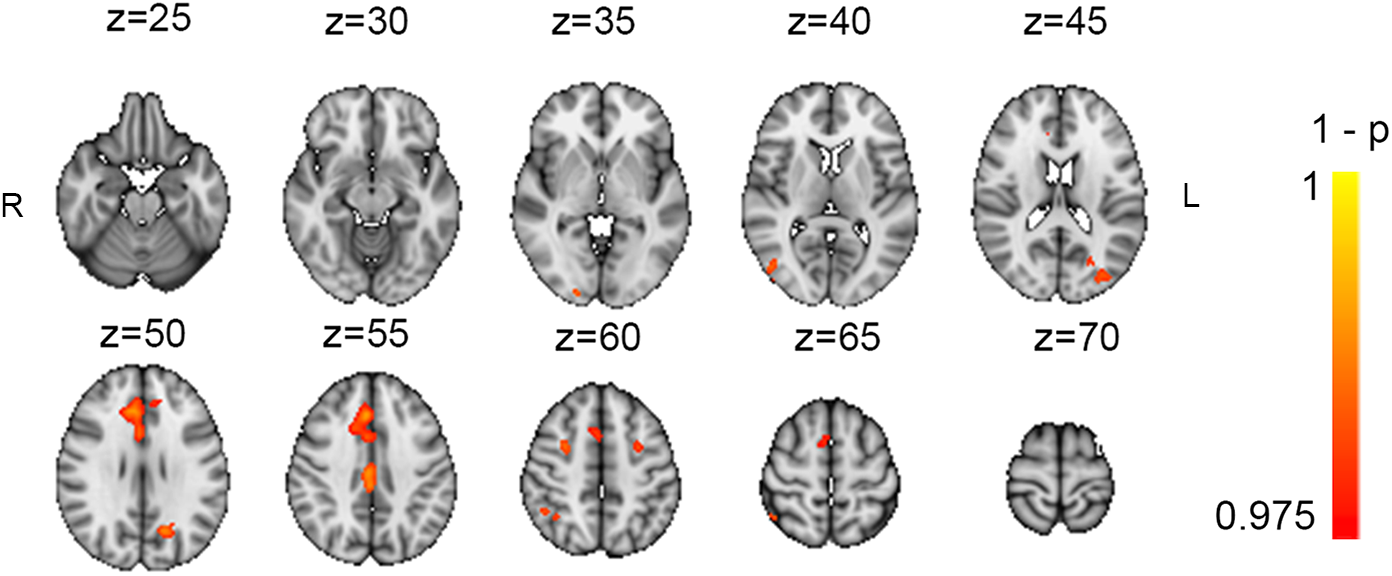
Association between Cogmed average performance score, and difference in task-related activation (L1 & L2 > PV) between timepoint 3 and 2.

### tDCS –fMRI associations

Results revealed no difference in fMRI-activation between the active and sham tDCS stimulation groups.

## Discussion

In this study we investigated the feasibility and effectiveness of combining CCT and tDCS, and tested the predictive value of and training-related changes in fMRI-based brain activation following an intervention with a commercially available working memory training package.

Tailoring behavioral interventions for alleviating cognitive difficulties following stroke calls for investigation of both treatment response, as well as delineating factors predicting favorable outcome. Cognitive abilities prior to training as well as motivation (Guye et al., 2017; Jaeggi et al., 2014) are currently among our best predictors for beneficial effects. Our results revealed significant improvement on all included tasks from the training program, in line with previous reports (Spencer-Smith and Klingberg, 2015; Westerberg et al., 2007). Our results support that repeated practice, where level of difficulty is adjusted during training, improves performance on the trained tasks. It is important to note that the current study investigates possible biomarkers for individual training outcome, and its beyond our aim to evaluate potential behavioral transfer effects. However, the generalizability of training-effects following CCT has been debated (Au et al., 2015; Melby-Lervåg and Hulme, 2013; Morrison and Chein, 2011; Shipstead et al., 2012; von Bastian and Oberauer, 2014). Indeed, recent large-scale reviews suggest a lack of reliable transfer beyond tasks that share properties with the trained tasks (Melby-Lervåg et al., 2016; Sala and Gobet, 2019). The lack of transfer effects is suggested to be caused by the failure of targeting common mechanisms underlying fluid intelligence such as working memory and attention, and rather targeting task specific mechanisms through repeated practice (Melby-Lervåg et al., 2016). Despite our promising behavioral results on trained tasks, the lack of transfer-effects reported in the literature calls for caution before implementation in a clinical setting.

Our results revealed no significant interaction between cognitive improvement and tDCS stimulation with the exception of one of the trained tasks, which did not reach significance after removal of an outlier. Although anodal tDCS has shown beneficial effects on cognitive training compared with sham stimulation in healthy adults (Martin et al., 2013) and have been associated with steeper learning curves (Ruf et al., 2017), a recent meta-analysis provided no support of a beneficial effect of tDCS on cognitive training compared to sham (Medina and Cason, 2017). The current literature investigating the added effect of tDCS when combined with working memory training in stroke patients is scarce. Single session anodal tDCS towards the left DLPFC improved recognition accuracy during an N-back task (Jo et al., 2009), and longitudinal concurrent anodal tDCS and CCT increased accuracy on a non-trained continuous performance test (CPT) compared to sham (Park et al., 2013). However, divergence in study protocols and results hampers a direct comparison (Filho et al., 2017). Beyond robust improvement on all trained tasks during the course of the intervention, our results provide no evidence of additional beneficial effects of tDCS stimulation. Intriguingly, current investigations aim to evaluate how home-based tDCS may aid patients with mild cognitive impairments (Park et al., 2019). This will potentially provide novel insights regarding clinical application for tDCS.

Our fMRI results for task activation during the first assessment revealed increased activation of the DAN, along with deactivation of DMN for passive viewing, load 1 and load 2. Further, contrasting tracking to passive viewing, and load 2 to load 1 revealed that attentive tracking increased DAN activation in a load-dependent manner, in line with previous reports based on healthy samples (Alnæs et al., 2014; Dørum et al., 2016).

Previous investigations suggest favorable reliability in task based compared to resting-state fMRI (Specht, 2020), indicating higher reliability when the task elicits strong patterns of activation (Johnstone et al., 2005) increasing contrast to noise ratio (Bennett and Miller, 2010). Our results are in line with these reports, as results revealed fair (>0.4) to good (>0.75) reliability across task-associated brain regions following guidelines for ICC interpretation (Cicchetti, 1994), with poor reliability in non-task associated areas.

Our results revealed no significant interaction between activation during passive viewing, load 1, load 2, tracking nor load, and time. Although we did not find any predictive value of fMRI-activation patterns on Cogmed performance improvement, our results revealed an association between average performance during the Cogmed training period and increased tracking-related activity in key nodes of the DAN and DMN as well as ACC. DMN is commonly suppressed during task engagement, with increasing suppression with increased task demands (Anticevic et al., 2012). This have been further supported by results indicating that individuals with lower cognitive abilities elicit a stronger deactivation of DMN during task engagement (Lipp et al., 2012), possibly indicating an increased effort to resolve the task.

In line with these findings our results indicate less suppression of PCC and precuneus during task engagement, with greater average performance.

Working memory capacity, reflecting both executive control as well as cognitive flexibility, has been linked to general cognitive abilities (Conway et al., 2003). Furthermore, higher cognitive abilities have been associated with increased activity in frontal and parietal cortices (Basten et al., 2015), and cognitive flexibility have been linked to brain activation in ventrolateral prefrontal cortex, as well as anterior and posterior cingulate (Dajani and Uddin, 2015). As our results indicate that greater cognitive capacity is associated with an increase in activation during task engagement over time, it may reflect compensation and learning to more efficiently resolve the task.

The current findings should be interpreted in light of several limitations. Based on the clinical ratings (NIHSS) at hospital discharge, the patients were sampled from the less severe part of the stroke severity spectrum. Further, patient drop-out during the intervention was relatively high, partly due to the labor-intensity of the training. Hence, our sample is biased towards higher functioning stroke patients, rendering the generalizability of our results to more severe cases unclear. The lack of a control group for the CCT does not allow us to disentangle effects of the cognitive training from the effect of time, anticipation, or other confounders. As our main aim was to identify potential fMRI-based markers for training outcome, we did not assess possible cognitive and clinical transfer effects, which will be relevant for evaluating clinical relevance, and further studies are needed to establish the premises for optimizing the clinical potential for working memory interventions in individual stroke patients. It has furthermore been reported that potential beneficial effects of tDCS may emerge at a delayed stage after intervention (Goodwill et al., 2016; Li et al., 2019). As our last assessment was performed shortly after the final stimulation, any delayed effects may have been lost. Future follow-up assessments of the same patients may provide relevant information about potential long-term effects of CCT and tDCS. Potential effects also may be dependent of stimulation frequency (Boggio et al., 2007; Yun et al., 2015). Increasing frequency beyond current protocol may have altered our results. Lesion size and location may influence current flow and cortical excitability (Marquez et al., 2013), and future studies may test for associations between lesion characteristics and response to tDCS. While the degree of cognitive impairment at baseline is likely to influence the response to cognitive training and rehabilitation in general, we found no significant association between baseline cognitive performance and training gain. However, the lack of pre-stroke neuropsychological evaluation complicates the assessment of stroke-related cognitive impairment. The group level task activation patterns in the current study largely mirrored those reported in previous studies in healthy controls (Alnæs et al., 2014; Dørum et al., 2016). However, the current study design did not allow us to make an explicit comparison (D’Esposito et al., 2003). Here we only considered brain activation in response to task demands. Future studies should also pursue a promising line of research utilizing imaging indices of structural and functional connectivity to investigate behavioral correlates of stroke, which may supplement and increase our understanding of stroke and stroke recovery.

In conclusion, we have investigated response to a computerized cognitive training program in 54 stroke survivors in combination with tDCS, as well as the predictive value of neural activation on training outcome, and neural alterations following training. Our results revealed increased performance across all trained tasks, with no additional benefit of tDCS. Brain activation prior to the training was not predictive for training outcome, nor was training gains reflected in altered brain activation. The generalizability of the reported beneficial effects of CCT remains uncertain, and future studies may be able to assess the transfer value of for relevant cognitive and clinical variables in stroke patients.

## Acknowledgements

This work was supported by the South-Eastern Norway Regional Health Authority (2014097, 2015044, 2015073), the Norwegian ExtraFoundation for Health and Rehabilitation (2015/FO5146), the Research Council of Norway (249795, 262372) and the European Research Council under the European Union’s Horizon 2020 research and Innovation program (ERC StG, Grant 802998).

## Notes

Conflict of Interest: We declare no competing financial interests. We obtained appropriate ethical approval from the local ethics committee, and all procedures were in line with the declaration of Helsinki. Relevant data have been made available at osf.io: https://osf.io/ndpuz/.

### Competing Interest Statement

The authors have declared no competing interest.

