## Supplementary material for "Reliability, sensitivity and predictive value of fMRI during multiple object tracking as a marker of cognitive training gain in combination with tDCS in stroke survivors"

Supplementary table 1:

| | Active stimulation group | | | | Sham stimulation group | | | |
|---|---|---|---|---|---|---|---|---|
| | Mean | sd | Min | Max | Mean | sd | Min | Max |
| **Participant descriptives (current)** | | | | | | | | |
| Age at inclusion | 69.78 | 7.68 | 48.08 | 80.12 | 69.54 | 7.31 | 47.61 | 81.88 |
| Males (%) | 62.96 | | | | 85.19 | | | |
| Time between stroke and inclusion (months) | 26.18 | 8.87 | 12 | 45 | 25.27 | 9.61 | 6 | 44 |
| Self-reported education in years | 14.08 | 3.07 | 9 | 20 | 14.69 | 4.27 | 9 | 30 |
| MMSE at inclusion | 28.19 | 1.47 | 25 | 30 | 27.78 | 2.17 | 22 | 30 |
| IQ | 109 | 15.95 | 84 | 135 | 111 | 18.11 | 66 | 136 |
| Days between inclusion and training | 35.26 | 14.62 | 18.96 | 70 | 32.29 | 11.19 | 20 | 74 |
| Days between MRI assessment pre/post training | 33.37 | 6.19 | 27 | 49 | 30.08 | 5.99 | 19 | 43 |
| **Clinical variables during hospitalization** | | | | | | | | |
| NIHSS at discharge | 1.5 | 1.62 | 0 | 7 | 1.17 | 1.46 | 0 | 6 |
| TOAST classification | Large vessel disease n=11<br>Small vessel disease n=8<br>Cardioembolic disease n=3<br>Other n=5 | | | | Large vessel disease n=9<br>Small vessel disease n=10<br>Cardioembolic disease n=3<br>Other n=5 | | | |

**Supplementary Table 1:** Sample characteristics, specified for the two separate intervention groups.

Supplementary table 2:

| | Grid | Sort | Digits | Cube | Hidden objects | Twist | 3D-Cube | Rotation |
|---|---|---|---|---|---|---|---|---|
| Session | **8.66** **(<0.001)** | **6.40** **(<0.001)** | **12.79** **(<0.001)** | **7.44** **(<0.001)** | **11.03** **(<0.001)** | **5.16** **(<0.001)** | **3.16** ***(0.002)*** | **2.21** ***(0.028)*** |
| Age | **-3.05** ***(0.004)*** | **-2.11** ***(0.041)*** | **-6.07** **(<0.001)** | **-2.54** ***(0.015)*** | **-4.86** **(<0.001)** | **-3.36** ***(0.002)*** | **-3.90** **(<0.001)** | **-3.41** ***(0.001)*** |
| Sex | -0.89 (0.377) | -0.33 (0.741) | -2.29 (0.027) | -1.22 (0.23) | -2.25 (0.029) | -1.25 (0.219) | -1.42 (0.162) | -1.34 (0.187) |
| tDCS | -0.65 (0.521) | -1.03 (0.309) | -0.06 (0.951) | -0.53 (0.599) | 0.12 (0.908) | 0.35 (0.73) | -0.82 (0.414) | -0.81 (0.422) |
| Education | 0.06 (0.951) | 0.96 (0.34) | 0.90 (0.374) | 0.14 (0.886) | 1.57 (0.124) | -0.12 (0.903) | -0.15 (0.884) | 0.33 (0.744) |
| Session x tDCS | 0.2 (0.842) | -0.39 (0.7) | -1.53 (0.128) | 0.46 (0.644) | -1.79 (0.074) | -1.76 (0.079) | 1.72 (0.087) | 1.23 (0.22) |

**Supplementary table 2**. Associations between training session, age, sex, tDCS, and educational level, and performance on Cogmed subtests t / (p), after removal of one outlier on the twist task. Raw p-vaules are displayed, significant findings are highlighted (FDR-corrected, alpha = 0.05)

Supplementary table 3:

| | CogMed change score | CogMed average performance score | CogMed change score outlier removed | CogMed average performance score outlier removed |
|---|---|---|---|---|
| Number of lesions | 0.07 (0.941) | 0.29 (0.774) | 0.31 (0.759) | 0.46 (0.648) |
| Global lesion volume | **-2.47** **(0.017)** | 0.35 (0.726) | -0.58 (0.563) | 0.61 (0.546) |
| Interaction | 1.01 (0.319) | -1.00 (0.321) | 0.63 (0.533) | -1.11 (0.271) |

**Supplementary table 3.** Statistical associations between the derived Cogmed scores, number of insults, lesion volume as well as the interaction between them. Results both with and without the significant outlier are displayed.

Supplementary table 4:

| | Direction | Size | x | y | z | Location |
|---|---|---|---|---|---|---|
| Frontal | pos | 110 | 30.5 | 64.3 | 61.3 | MFG |
| | pos | 65 | 56.8 | 63.7 | 61.2 | MFG |
| Limbic | pos | 1212 | 41.4 | 74.7 | 52.6 | ACC |
| | pos | 30 | 50.2 | 81.3 | 49.7 | ACC |
| | pos | 203 | 43.8 | 51.9 | 53.7 | PCC |

**Supplementary table 4.** Cogmed average score clusters associated with difference in activation between timepoint 2 and 3 for all included Cogmed tests. (all p<.05, corrected)

Supplementary table 5:

| | Correlation cabpad baseline and Cogmed change score | | | Interaction Cogmed change score and tdcs | |
|---|---|---|---|---|---|
| test | r | t | p | F | p |
| Fingertap right | -0.03 | -0.20 | 0.838 | 0.09 | 0.76 |
| Fingertap left | 0.00 | 0.01 | 0.994 | 0.53 | 0.47 |
| FAS fonetic flow | -0.03 | -0.22 | 0.826 | 0.11 | 0.75 |
| FAS semantic flow | -0.05 | -0.37 | 0.713 | 2.31 | 0.13 |
| Stroop Number of responses | 0.00 | 0.03 | 0.975 | 1.11 | 0.30 |
| Spatial memory | 0.00 | 0.00 | 0.999 | 0.26 | 0.61 |
| Coding correct responses | -0.11 | -0.80 | 0.427 | 0.42 | 0.52 |

**Supplementary table 5:** Summary statistics of association between cognitive performance at baseline (TP1) measured with the Cabpad battery, Cogmed change score, and tDCS. As results revealed no significant results, raw uncorrected statistics are displayed.

Supplementary figure 1:

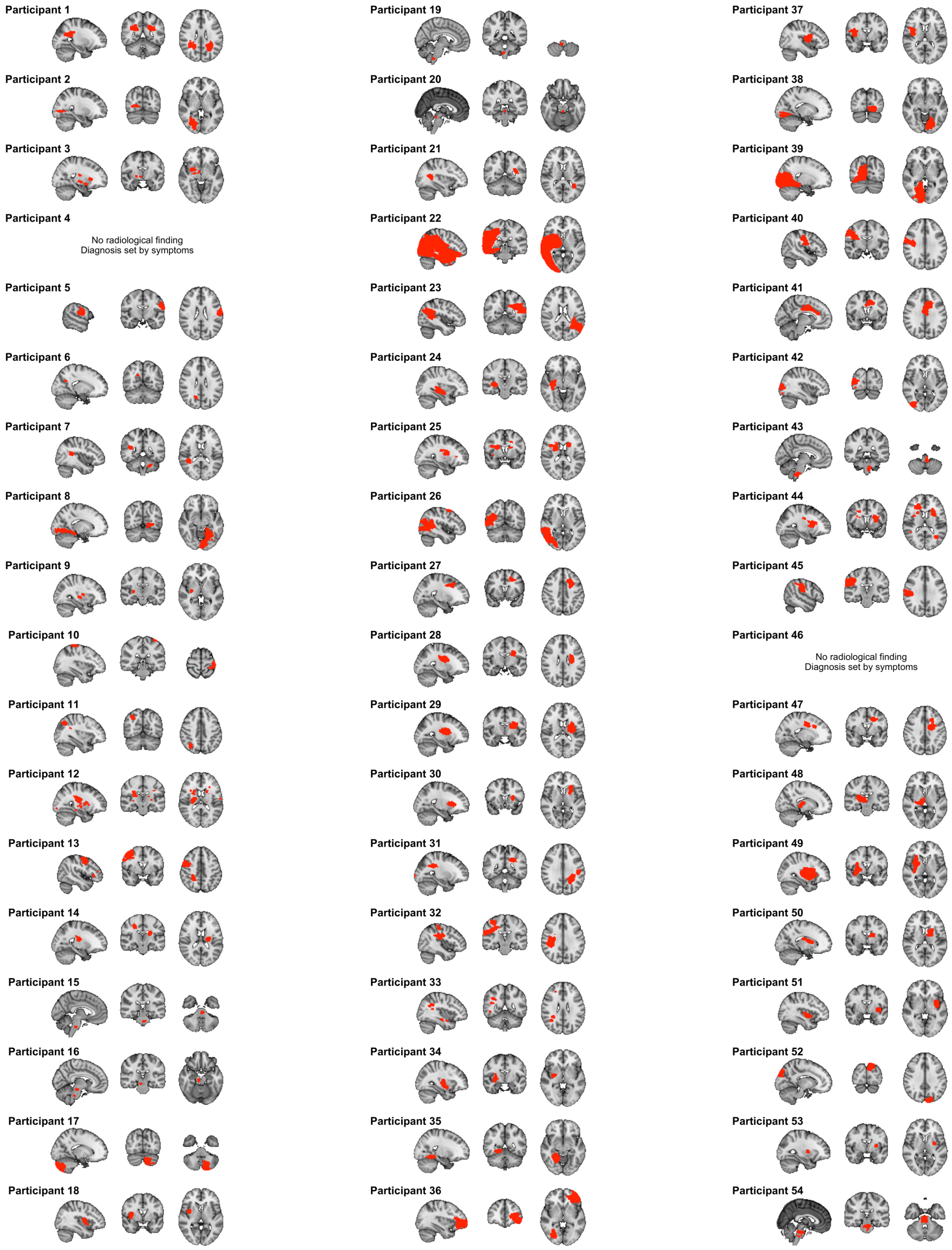

**Supplementary figure 1.** Graphical display of lesion location for participants completing the intervention.

Supplementary figure 2:

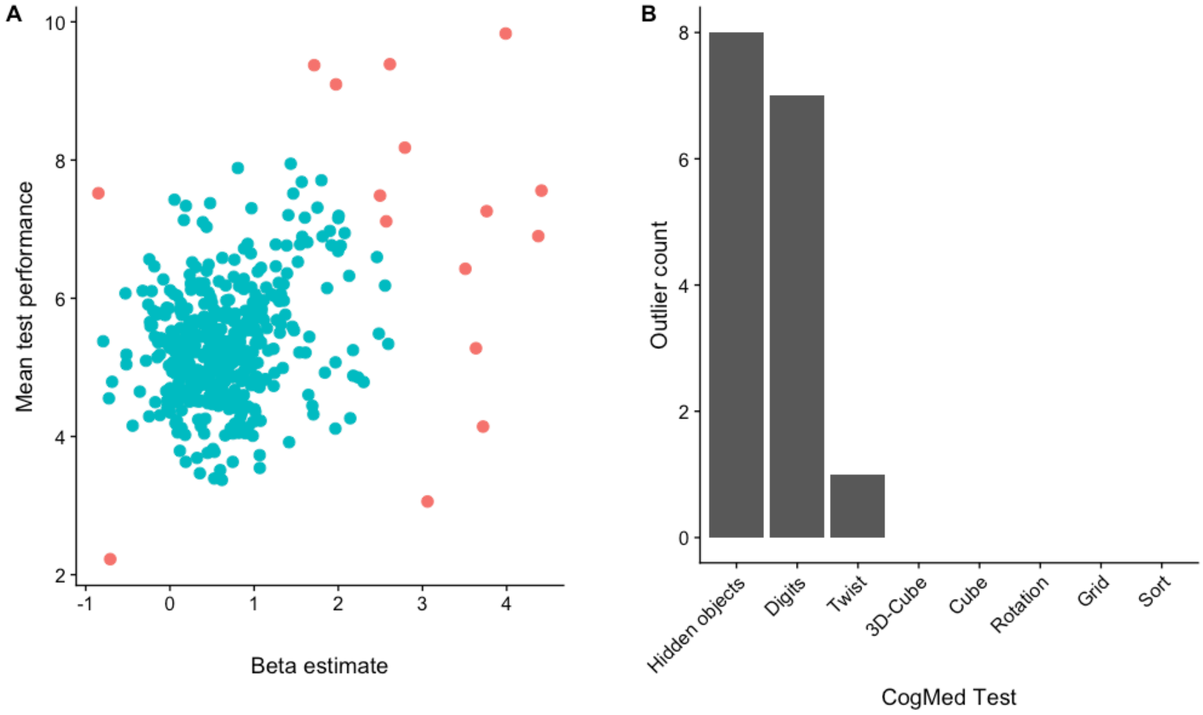

**Supplementary figure 2.** Overview over (A) distribution of values from bivariate outlier detection where outliers marked in red, and (B) count of outliers for each subtest separately.

Supplementary figure 3:

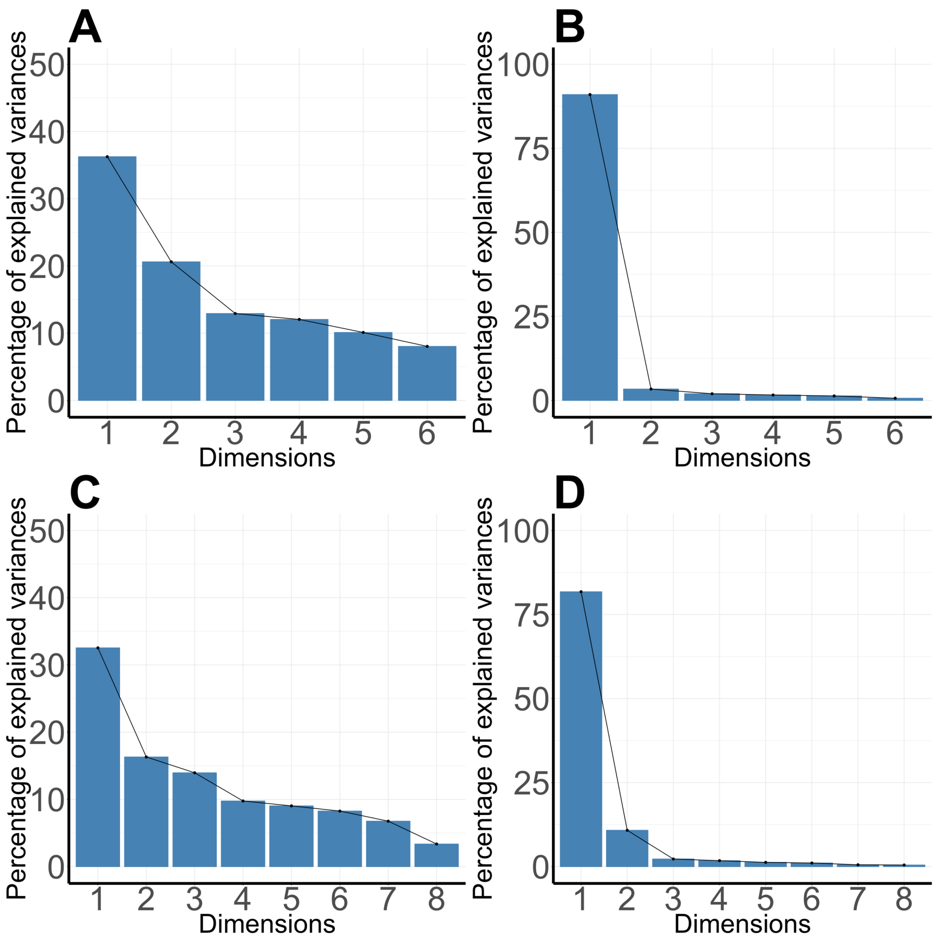

**Supplementary figure 3**. Scree-plot displaying how Cogmed scores loads on PCA factors, for (A): Beta-estimates from the six subtests included in further analysis, and (B): Mean performance across the six subtests included in further analysis and (C): Beta-estimates and (D): Mean performance across with the Cogmed-tests identified as outliers included.

Supplementary figure 4:

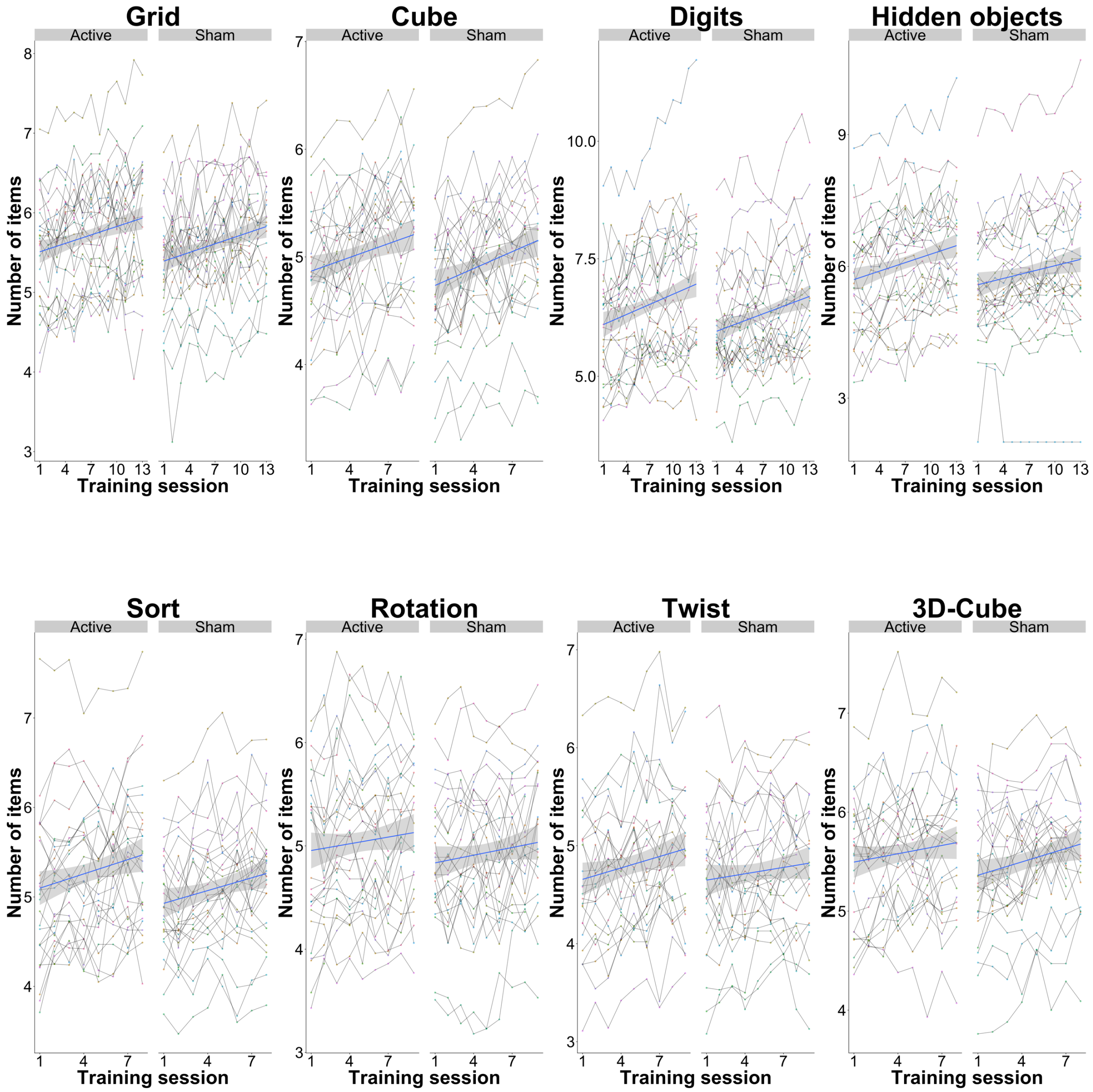

**Supplementary figure 4.** Overview over association between time and performance during the Cogmed training-period, with the outlier in the twist task removed.

Supplementary figure 5:

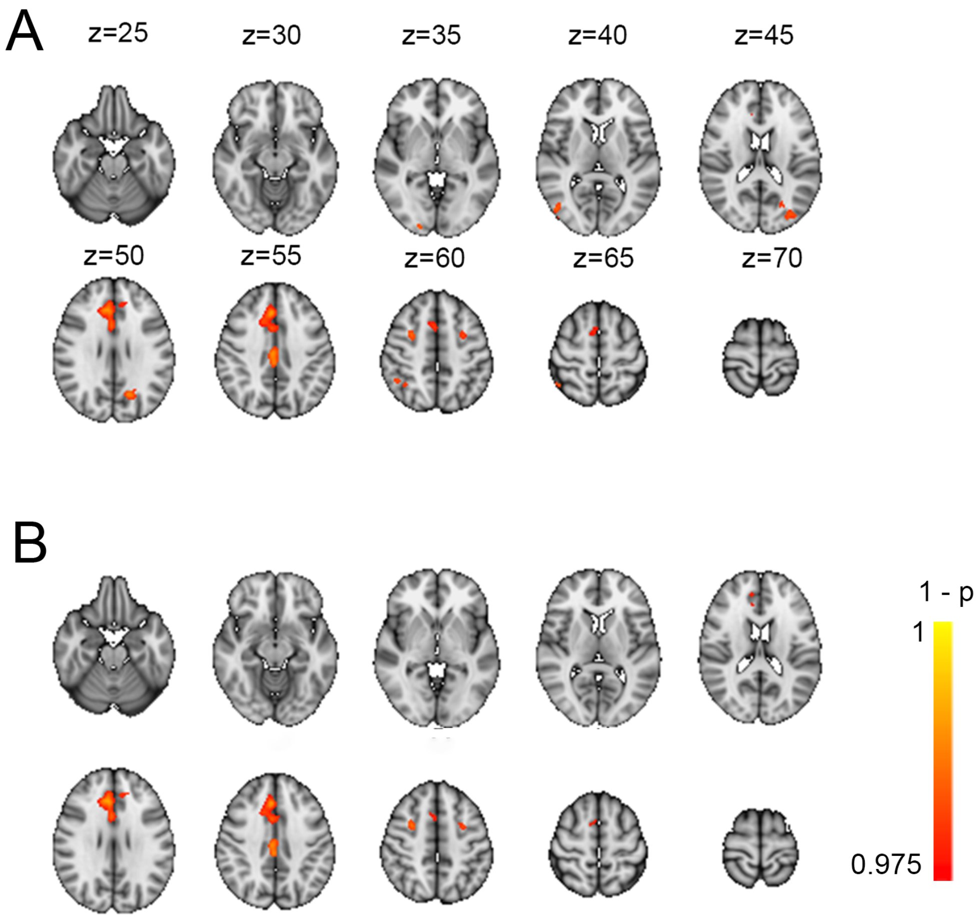

**Supplementary figure 5**. Comparison of results from randomise, comparing Cogmed average performance score based on (A): six subtests, and (B): eight subtests. Corresponding t-maps revealed a correlation of 0.99.

Supplementary figure 6.

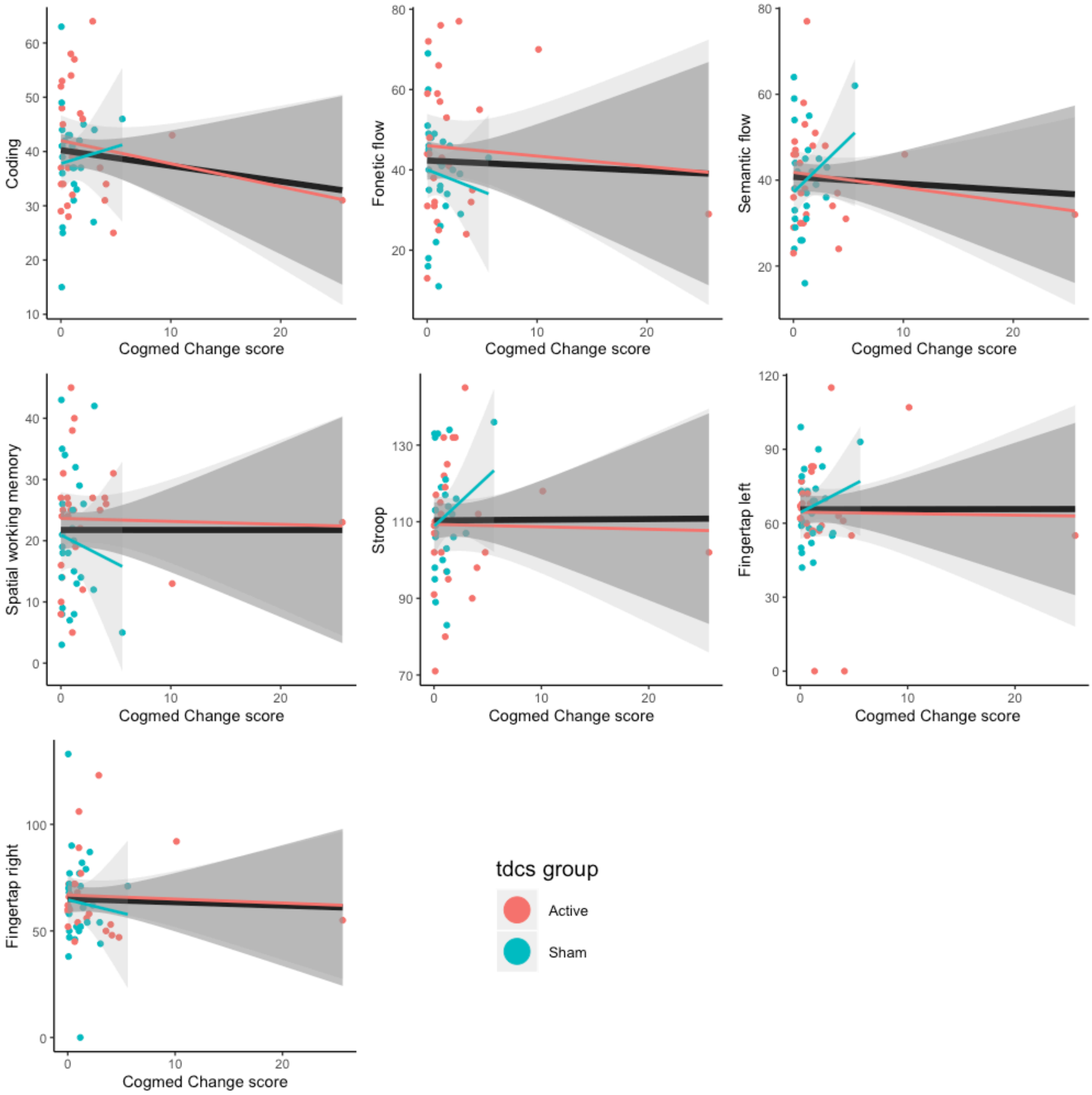

**Supplementary figure 6:** Association between cognitive testing prior to training, and Cogmed performance score. Supplementary table 4 provides detailed statistics.

Supplementary figure 7.

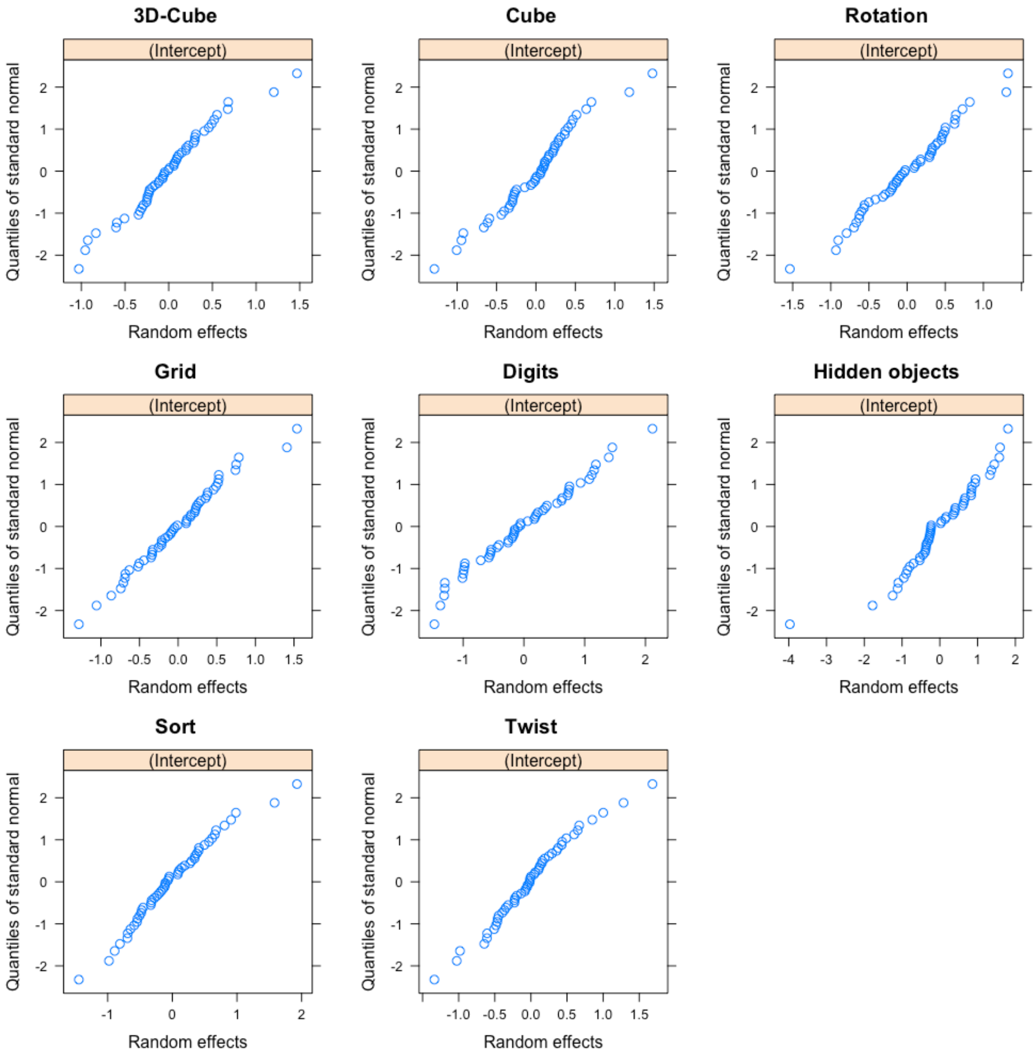

**Supplentary figure 7:** QQ-plot displaying distribution of intercepts of random effects of the lme. Visual inspection did not indicate severe violations of the criteria.
